# Time-varying transmission dynamics of Novel Coronavirus Pneumonia in China

**DOI:** 10.1101/2020.01.25.919787

**Authors:** Tao Liu, Jianxiong Hu, Jianpeng Xiao, Guanhao He, Min Kang, Zuhua Rong, Lifeng Lin, Haojie Zhong, Qiong Huang, Aiping Deng, Weilin Zeng, Xiaohua Tan, Siqing Zeng, Zhihua Zhu, Jiansen Li, Dexin Gong, Donghua Wan, Shaowei Chen, Lingchuan Guo, Yan Li, Limei Sun, Wenjia Liang, Tie Song, Jianfeng He, Wenjun Ma

## Abstract

**Rationale:** Several studies have estimated basic production number of novel coronavirus pneumonia (NCP). However, the time-varying transmission dynamics of NCP during the outbreak remain unclear.

**Objectives:** We aimed to estimate the basic and time-varying transmission dynamics of NCP across China, and compared them with SARS.

**Methods:** Data on NCP cases by February 7, 2020 were collected from epidemiological investigations or official websites. Data on severe acute respiratory syndrome (SARS) cases in Guangdong Province, Beijing and Hong Kong during 2002-2003 were also obtained. We estimated the doubling time, basic reproduction number (*R_0_*) and time-varying reproduction number (*R_t_*) of NCP and SARS.

**Measurements and main results:** As of February 7, 2020, 34,598 NCP cases were identified in China, and daily confirmed cases decreased after February 4. The doubling time of NCP nationwide was 2.4 days which was shorter than that of SARS in Guangdong (14.3 days), Hong Kong (5.7 days) and Beijing (12.4 days). The *R_0_* of NCP cases nationwide and in Wuhan were 4.5 and 4.4 respectively, which were higher than *R_0_* of SARS in Guangdong (*R_0_*=2.3), Hongkong (*R_0_*=2.3), and Beijing (*R_0_*=2.6). The *R_t_* for NCP continuously decreased especially after January 16 nationwide and in Wuhan. The *R_0_* for secondary NCP cases in Guangdong was 0.6, and the *R_t_* values were less than 1 during the epidemic.

**Conclusions:** NCP may have a higher transmissibility than SARS, and the efforts of containing the outbreak are effective. However, the efforts are needed to persist in for reducing time-varying reproduction number below one.

**At a Glance Commentary:** *Scientific Knowledge on the Subject:* Since December 29, 2019, pneumonia infection with 2019-nCoV, now named as Novel Coronavirus Pneumonia (NCP), occurred in Wuhan, Hubei Province, China. The disease has rapidly spread from Wuhan to other areas. As a novel virus, the time-varying transmission dynamics of NCP remain unclear, and it is also important to compare it with SARS.

*What This Study Adds to the Field:* We compared the transmission dynamics of NCP with SARS, and found that NCP has a higher transmissibility than SARS. Time-varying production number indicates that rigorous control measures taken by governments are effective across China, and persistent efforts are needed to be taken for reducing instantaneous reproduction number below one.

## Introduction

Eighteen years ago, severe acute respiratory syndrome (SARS) broke out globally, which caused more than 8000 cases with a fatality rate of 9.6% (1). Since December 2019, a pneumonia infection with 2019-nCoV, which was now named as Novel Coronavirus Pneumonia (NCP), broke out in Wuhan, Hubei Province, China, and rapidly spread throughout China and to many other countries (2–4). As of February 9, 2020, 2019-nCoV had transmitted to 34 provinces, regions and municipal cities across China. A total of 40,235 confirmed NCP cases with 908 deaths (2.3%) and 6,484 (16.1%) critically ill cases, and there were still 23,589 suspected cases (5).

As an emerging infectious disease, we need to quickly understand its etiological, epidemiological and clinical characteristics to take prevention and control measures. Several studies have described the epidemiological and clinical characteristics of NCP, and have demonstrated that it can transmitted between humans (4, 6). A few studies have also estimated the basic reproduction number (*R_0_*) of NCP (6–11). For instance, Li et al. computed the *R_0_* of 2.2 using daily NCP data before January 4 in Wuhan. Our study using early epidemic stage data also found reproductive number of NCP was 2.9 (11). However, most of these studies employed daily reporting cases at very early epidemic stage or were purely based on mathematical modeling. It has been suggested that epidemic size may be seriously under-reported due to the unknown etiology and lack of diagnosis protocol at early epidemic stage(12), which may lead to underestimation of *R_0_*. In addition, for control measures to be optimized during ongoing NCP epidemics, temporal changes in reproduction number must be tracked. However, none of previous studies have estimated time-varying instantaneous reproduction number (*R_t_*), which is an important parameter for assessing whether the control and prevention measures are effective or whether additional measures are needed (13). Moreover, excerpt for Hubei Province, most of confirmed cases reporting from other provinces were imported from Hubei Province at early epidemic stage. With increasing secondary cases in those provinces, however, it is unknown how much the *R_t_* for secondary cases is in those Provinces. Knowing this information is very helpful to assess the epidemic trend and the effects of control measures in those provinces.

As a milestone of global public health events, the transmission dynamics of SARS have been well studied (14, 15). Comparing the transmissibility of NCP with SRAS could imply decision making on the prevention and control of NCP. Therefore, in this study, we described the epidemiological characteristics of NCP using the latest data across China, and estimated the basic and time-varying reproduction numbers which were compared with SARS. Our findings are critical to optimize prevention and control measures during the ongoing epidemic of NCP, as well as guide clinical management.

## Methods

### Data Collection

Daily number of confirmed NCP cases across China and in other countries were collected as of February 7, 2020. Confirmed NCP case was defined based on the Diagnosis and Treatment Scheme of NCP released by the National Health Commission of China (Section 1.1 in Supplementary materials) (16). For cases in Guangdong Province, China, individual data including date of onset, hospitalization and diagnosis were obtained from medical records and epidemiological investigations. We collected daily number of reported cases in other provinces of China or other countries from official websites.

We obtained daily number of SARS cases during 2002-2003 in Guangdong Province from Guangdong Provincial Center for Disease Control and Prevention (GDCDC). In addition, daily number of SARS cases in Beijing (from March 5 to May 29, 2003) and Hong Kong (from February 15 to May 31, 2003) were obtained from Pang et al.’s report (17) and WHO website (18), respectively.

### Estimating and nowcasting the daily incidence of NCP

Because only the daily number of reporting cases were obtained from regions out of Guangdong Province, we estimated the daily number of incidences using a generalized additive model (GAM) (Section 1.2 in Supplementary materials). First, we collected each individual onset date and reporting date in Guangdong province which was treated as a sample of all confirmed cases nationwide. Second, a GAM model was used to establish the relationship between onset date and reporting date, and obtained the lagged probability distribution of daily number of incidences for the number of reporting cases. Third, we used the lagged probability distribution to predict the daily number of incidences based on the daily number of reporting cases in other provinces of China, which was added up to obtain the daily incidences nationwide. In order to test the validity of this approach, we randomly divided all cases in Guangdong into two groups (group A and B). Cases in group A was used to establish a GAM model, which was then used to predict the number of incidences in group B. We then estimated the correlation between the actual incidence and the estimated incidence, and the results of cross-validation showed that the R^2^ was 89.7%, and the root mean square error (RMSE) was 4.11 (Figure S1).

The number of occurred-but-not-yet-reported cases were further estimated during the study period, which caused by reporting delays. Reasons for such delays include the time to complete diagnosis test, logistics, and overwhelmed surveillance systems. To minimize the effects of reporting delay, we applied a Bayesian nowcasting model (Nowcasting by Bayesian Smoothing) to complement the daily total number of incidences (19). This model is characterized by its ability to deal with the uncertainty in the delay distribution and the time evolution of epidemic curve.

### Estimation of the reproduction number

#### Basic reproduction number (*R_0_*)

In the early phase of an infectious disease outbreak, the initial reproductive rate approximated an exponential growth rate (20). We selected a period of exponential growth in the epidemic curve to calculated *R_0_*, which was defined as the expected number of secondary cases produced by a typical primary case. Since the daily incidence data is an integer, Poisson regression was used to fit the exponential growth rate. We used a prior Gamma distribution for the generation time, with a shape parameter (mean generation time of 7.5 days) and a scale parameter (standard deviation of 3.4 days) from a previous study (7). We estimated *R_0_* of all confirmed NCP cases nationwide and in Wuhan, and *R_0_* of SARS cases in Guangdong Province, Hong Kong and Beijing. We also estimated *R_0_* for secondary NCP cases in Guangdong Province.

#### Time-varying instantaneous reproduction number (*R_t_*)

In contrast to *R_0_*, time-varying instantaneous reproduction number (*R_t_*) represents the average number of secondary cases that would be produced by a typical primary case infected at time *t* if conditions remained constant after time *t* (13). We estimated *R_t_* from the time series of onset cases, with the same Gamma distribution of generation time as *R_0_*. In order to maintain the accuracy of the prediction and without hiding the underlying time trend, *R_t_* values were estimated over a 10-day moving window. For the period of estimation, the end date was selected as the latest date of the available data, and the starting date was selected as the earliest date for which the 10-day instantaneous reproduction number estimated above could be assumed constant. We estimated *R_t_* for all confirmed NCP cases nationwide, and in Wuhan, and for secondary NCP cases in Guangdong Province.

#### Statistical analysis

We applied frequency and percentages (%) to describe categorical variables, and used mean±*SD* to describe the continuous variables. The doubling time of NCP and SARS were defined and estimated using a method proposed by Galvani et al (21).

#### Sensitivity analysis

A series of sensitivity analyses were conducted to quantify the effect of parameters changes on *R_0_* value. We changed the shape parameter of Gamma distribution from 7.0 to 8.0 days for estimating the *R_0_* of NCP cases, and changed the same parameter from 8.0 to 9.0 for estimating the *R_0_* of SARS cases.

R software (version 3.6.0) was used for data analyses, with “NobBs” package for building nowcast model, “*R_0_*” for calculating *R_0_* and “EpiEstim” for estimating *R_t_*. Two tailed *P*<0.05 were considered statistically significant for all statistical tests.

#### Ethics statement

Data collection and analysis of cases were determined by the National Health Commission of the People’s Republic of China to be part of a continuing public health outbreak investigation and were thus considered exempt from institutional review board approval.

## Results

### Description of the outbreak

As of February 7, 2020, a total of 34,598 confirmed cases were identified in 34 provinces, regions and municipal cities across China, and 270 confirmed cases were identified in other 24 countries (Figure 1A). Hubei Province reported the most cases (n=24,953), in which 13,603 (54.5%) cases were reported in Wuhan, the capital city of Hubei. The fatality rate of NCP was 2.8% (699/24,953) in Hubei Province, and 0.24 % (23/9,645) in other provinces, China. A total 1,075 confirmed cases were reported in Guangdong Province, in which 220 (20.5%) cases were secondary cases, 51 cases were identified with positive of 2019-nCoV but did not report any symptoms, and 2 secondary cases were infected by sharing elevator with the index cases from their neighbors (Figure 1B).

**Figure 1.**
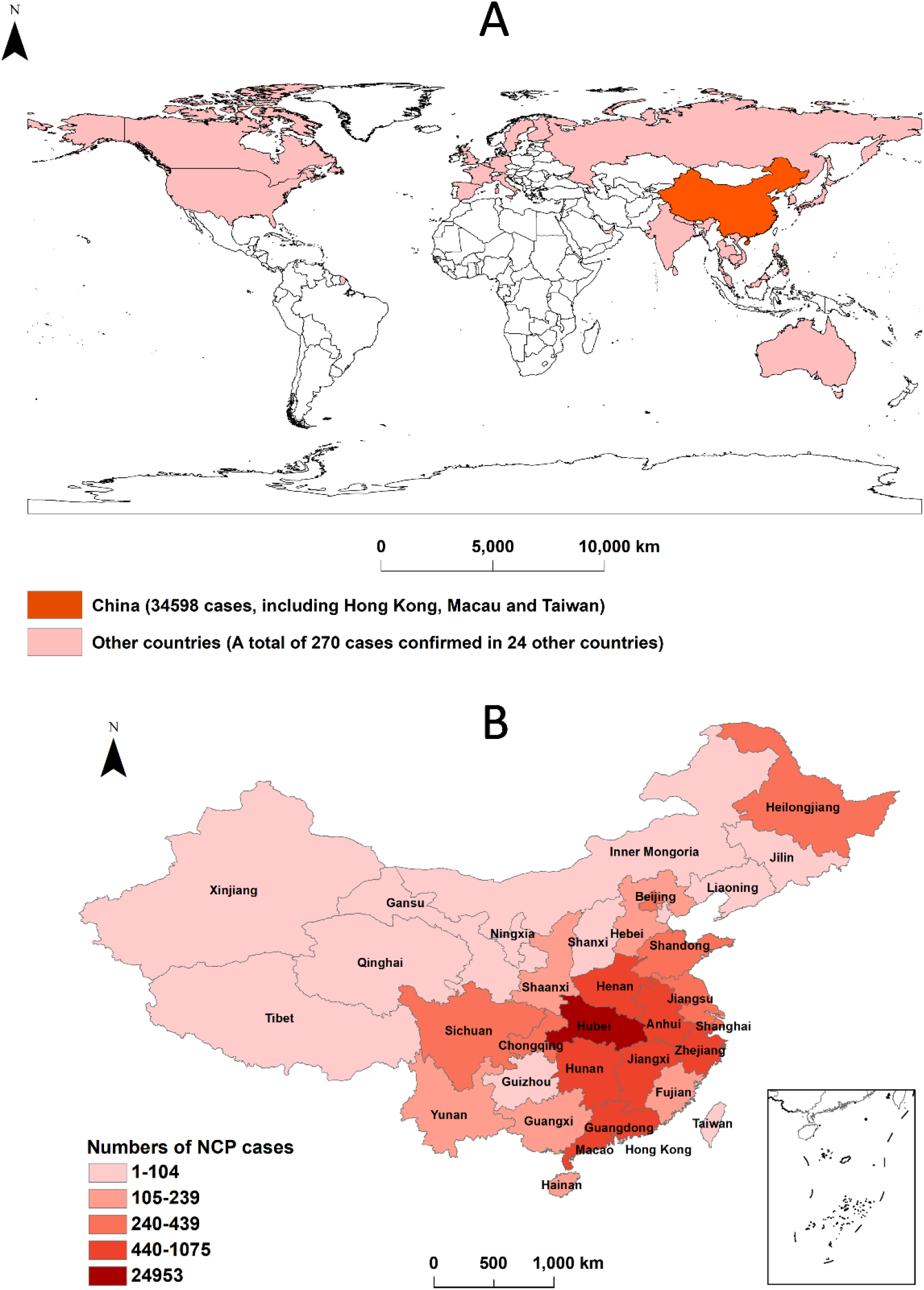
Distribution of NCP cases worldwide and in China, as of February 7, 2020. Panel A: The spatial distribution of NCP cases worldwide. Panel B: The spatial distribution of NCP cases in China.

Figure 2 displays the epidemic trend of NCP nationwide, in Wuhan and in Guangdong Province. Daily numbers of reporting cases continuously increased to the peak on February 4, 2020, and slightly decreased then after nationwide and in Wuhan. The peak of daily reporting cases was on January 31, 2020 in Guangdong Province, and then significantly decreased. The daily incidence after nowcasting showed similar trend with the reporting data. The doubling time of NCP was 2.4, 2.8, and 3.6 days nationwide, in Wuhan, and in Guangdong Province, respectively (Table 1).

**Figure 2.**
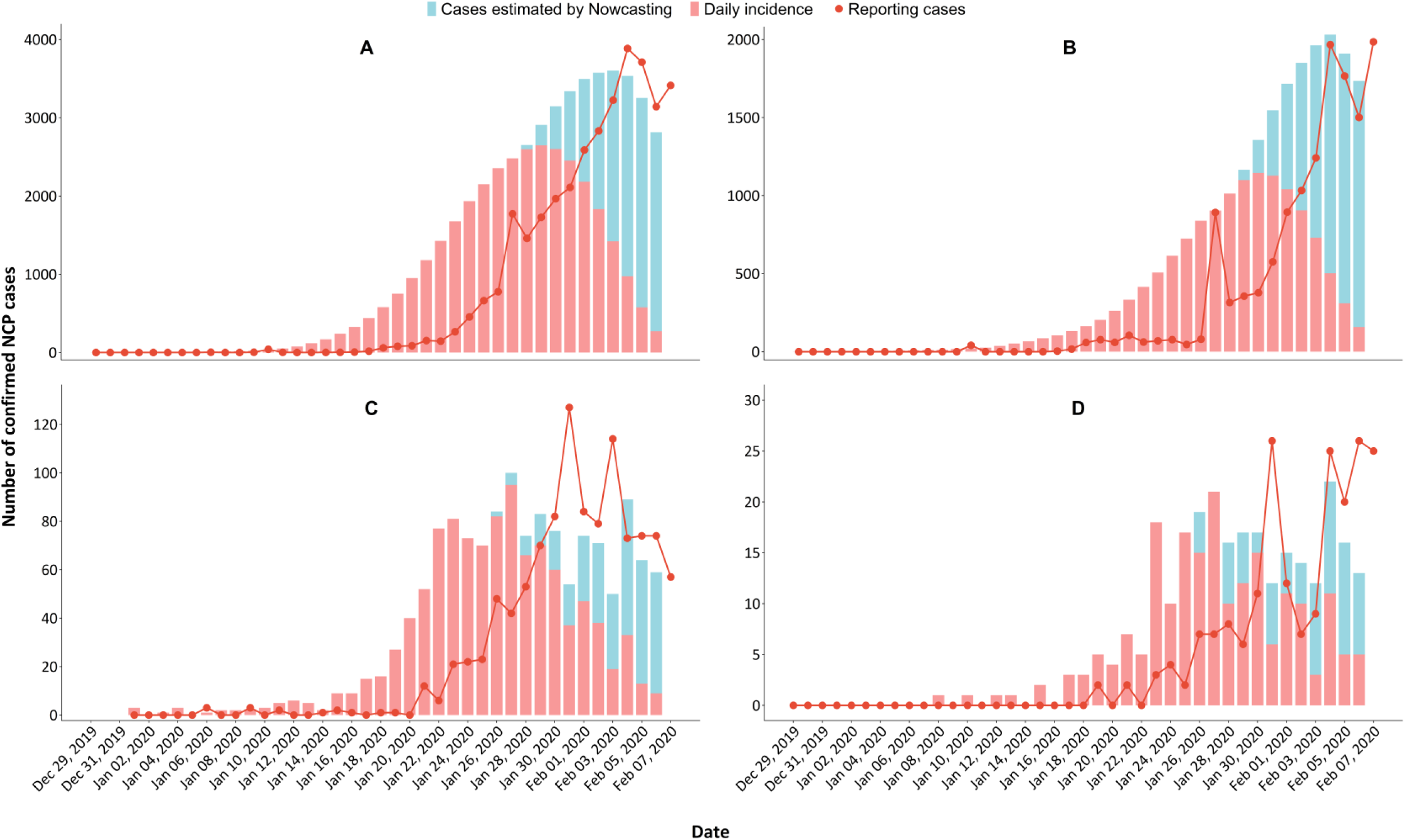
Temporal distribution of confirmed NCP cases nationwide, in Wuhan and in Guangdong Province. Panel A: Temporal distribution of NCP cases nationwide; Panel B: Temporal distribution of NCP cases in Wuhan; Panel C: Temporal distribution of all confirmed NCP cases in Guangdong Province; Panel D: Temporal distribution of secondary NCP cases in Guangdong Province.

**Table 1.**
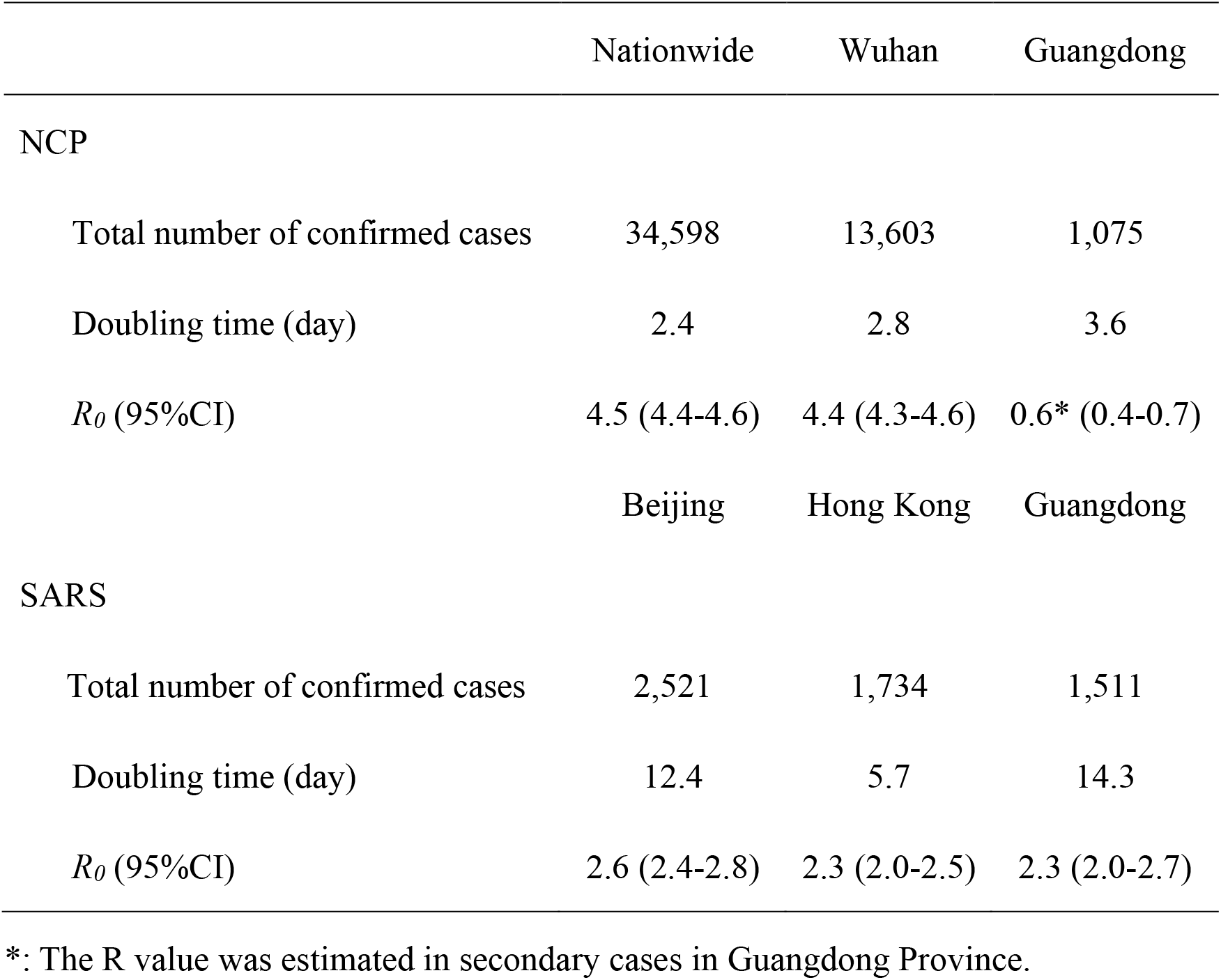
General characteristics of NCP cases nationwide, in Wuhan and in Guangdong Province, up to February 7, 2020

A total of 1,511, 1,734 and 2,521 SARS cases were reported in Guangdong Province, Hong Kong and Beijing during 2002-2003. The peak of daily cases in the three regions were found on February 8, March 25 and April 25, 2003 respectively (Figure S2). The doubling time of SARS in Guangdong, Hong Kong and Beijing were 14.3, 5.7 and 12.4 days during the period prior to the peak, respectively (Table 1).

### Estimation of basic reproduction number (*R_0_*)

The *R_0_* values of NCP were 4.5 (95%CI: 4.4-4.6) and 4.4 (95%CI: 4.3-4.6) nationwide and in Wuhan, respectively. The *R_0_* for secondary NCP cases in Guangdong Province was 0.6 (95%CI: 0.4-0.7). The *R_0_* values for SARS were 2.3 (95%CI: 2.0-2.7), 2.3 (95%CI: 2.0-2.5) and 2.6 (95%CI: 2.4-2.8) in Guangdong, Hong Kong and Beijing, respectively (Table 1).

### Estimation of time-varying reproduction number (*R_t_*)

During the epidemic, at national level, *R_t_* values increased from 6.9 (95%CI: 5.5-8.4) before January 9 to a peak of 8.8 (95%CI: 8.3-9.4) before January 16, and then continuously decreased to 1.59 (95%CI: 1.57-1.61) before February 6. The *R_t_* values in Wuhan continuously decreased from 6.2 (95%CI: 4.9-7.6) by January 9 to 2.10 (95%CI: 2.07-2.14) before February 6. By contrast, *R_t_* values for secondary NCP cases in Guangdong Province was kept at a low-level ranging from 0.8 (95%CI: 0.6-1.1) to 0.2 (95%CI: 0.2-0.3) during the epidemic (Figure 3).

**Figure 3.**
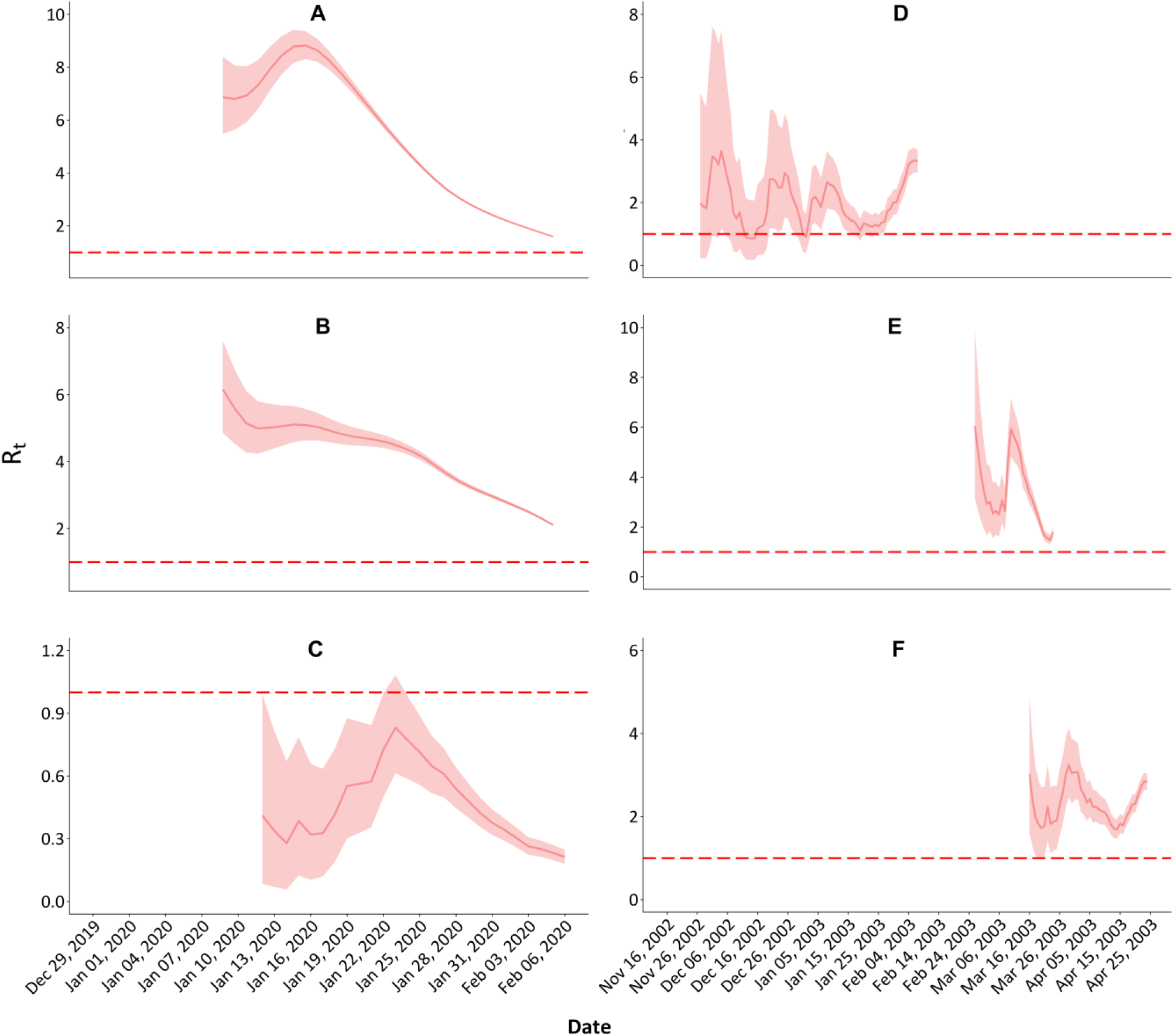
Time-varying reproduction number (*R_t_*) of NCP and SARS in China. Panel A: *R_t_* of NCP nationwide; Panel B: *R_t_* of NCP in Wuhan; Panel C: *R_t_* of secondary NCP in Guangdong Province; Panel D: *R_t_* of SARS in Guangdong Province; Panel E: *R_t_* of SARS in Hong Kong; Panel F: *R_t_* of SARS in Beijing;

### Sensitivity analyses

The results of sensitivity analyses showed the *R_0_* values of NCP and SARS were robust to the changes of GT (Figure S3). For example, the *R_0_* of NCP cases nationwide changed from 4.0 (95%CI: 3.9-4.1) using GT=7.0 to 4.9 (95%CI:4.8-5.1) using GT=8.0.

## Discussion

The 2019-nCoV is a novel coronavirus, which is different from SARS and other SARS-like viruses(3). However the time-varying transmission dynamics of NCP during the epidemic remain unclear(22). In this study, we estimated the *R_0_* of NCP infected with 2019-nCoV nationwide, and found that the *R_0_* was 4.5, which means an average of 4.5 secondary cases generated by a primary case in the early stage. An approaching *R_0_* of 4.4 was found in Wuhan. Our estimated *R_0_* was larger than that (*R_0_*=2.2) reported by Li et al. who employed the data in Wuhan between December 10, 2019 and January 4, 2020, the very early stage of this epidemic (7). It was debated that the number of NCP cases might be seriously underreported in the early stage (10, 23), since 2019-nCoV was confirmed on January 7, and the official diagnosis protocol was released by WHO until January 17 (12). In addition, the surveillance was not widely conducted, and many infections may be missed. Read et al. estimated that only 5.1% of infections in Wuhan were identified in the early period (23). Zhao et al. estimated that the *R_0_* of NCP was larger than 5.0 using national reported data from January 10 to January 24, 2020, under the scenario that the reporting rate was not changed during that period(12).

Although the fatality rate of NCP cases was much lower than SARS(1), we observed higher *R_0_* and much shorter doubling time of NCP than SARS (14, 24, 25), indicating the higher transmissibility of 2019-nCoV. These differences can be due to two reasons. First, epidemiological and clinical evidences suggest that asymptomatic or mild NCP cases during their incubation periods could effectively transmit 2019-nCoV, which was different from SARS because most SARS cases were infected by “super spreaders”, and SARS cases in incubation period and mild cases could not infected susceptible (26). The average incubation period of NCP was 4.8 days, ranging from 1 to 14 days, and the average period from onset to isolation was 2.9 days (11), which indicates a long transmission period of NCP. Second, NCP epidemic coincided with the approaching of Chinese Lunar New Year holiday, during which, under normal circumstances, an estimated 3 billion trips would be made, and 15 million trips occurred in Wuhan (27), which dramatically increased population mobility and accelerated the spread of NCP. Although Wuhan prohibited all transport in and out of the city as of 10:00 on January 23 2020, millions of citizens have left Wuhan before that time, and they may have become the major exporter of 2019-nCoV infection in other regions in China and other counties. Fortunately, the intervention of travel restriction was effective in slowing the 2019-nCoV invasion of new locations (28).

We observed significant decrease in the time-varying *R_t_* values of NCP nationwide and in Wuhan particularly after January 16. More importantly, the daily reporting cases began to decline after February 4. These results indicate that the rigorous measures of prevention and control taken by Chinese governments are taking into effect (Figure S4). For example, following the Wuhan travel restriction, most provinces and cities also implemented travel restriction, and conducted quarantine on all outpatients in fever, close contracts, and especially in persons travelling from Hubei Province. The Chinese Lunar New Year holiday was extended, and all gathering and public activities were restricted to prevent the transmission of 2019-nCoV. More importantly, almost all provinces and regions in China have initiated the highest level of public health emergency response. Risk communication and health education on NCP were broadly conducted through various channels, which has improved the general public to understand the risk of NCP, and take voluntary actions to detect, diagnose and treat cases infected 2019-nCoV earlier. This can be verified by decreasing period from onset to diagnosis during the epidemic in Guangdong Province (Figure S5).

We also observed that the *R_0_* was less than 1, and the *R_t_* values continuously declined after January 23 for secondary NCP cases in Guangdong Province, which demonstrates that the implemented prevention and control measures have significantly reduced the risk of indigenous outbreak of NCP, and that the increasing NCP cases were dominated by imported cases. Meanwhile, we found that two secondary cases were infected through sharing the public elevator with cases from their neighbors, which indicates the higher risk of transmission of NCP in community. Similar cases were also reported in other regions. For example, Wang et al. reported that 57 (including 40 healthcare workers and 17 patients) out of 138 hospitalized patients were presumed to have been infected with 2019-nCoV in a Wuhan hospital (29). Therefore, more efforts are needed to prevent the transmission in communities and other public places.

This study has several strengths. First, this is the first study, to date, to assess the time-varying transmission dynamics of NCP based on incidence data nationwide. We also estimated instantaneous reproduction number for secondary cases in Guangdong Province. Secondly, we compared the transmission dynamics of NCP with SARS using the same methods. Third, we initially assessed the effects of control measures previously taken by the governments. These findings could provide important information for optimizing control measures.

Our study has several limitations. First, the incidence date of confirmed cases in provinces except for Guangdong Province were estimated by a GAM modeling, which may lead to misclassification bias. However, the cross-validation test indicated a good performance of our GAM modeling. In addition, the *R_0_* (3.8, 95%CI: 3.4-4.2) estimated for imported cases in Guangdong Province, as a sample of the total confirmed cases nationwide, was comparable with the *R_0_* estimated nationwide, indicating that our results are robust. Second, we are still in the early stage of this outbreak, and there is much uncertainty in epidemiological parameters that affects transmissibility of NCP. Sequential studies are urgently needed to fill-in these knowledge gaps.

In summary, NCP may have a higher pandemic risk than SARS in 2003, and the efforts of containing the outbreak are taking into effect. Our findings indicate that rigorous control and prevention measures taken by Chinese governments on early detection, diagnosis and treatment of NCP cases are still needed to uphold until the time-varying reproduction number below one.

## Data sharing

The data that support the findings of this study are available from the corresponding author on reasonable request. Participant data without names and identifiers will be made available after approval from the corresponding author. After publication of study findings, the data will be available for others to request. The research team will provide an email address for communication once the data are approved to be shared with others. The proposal with detailed description of study objectives and statistical analysis plan will be needed for evaluation of the reasonability to request for our data. The corresponding author will make a decision based on these materials. Additional materials may also be required during the process.

## Acknowledgements

We thank all the medical and nursing staff who assisted in the care of patients; the members from health department and CDC in Guangdong Province for their contribution in data collection, 2019-nCoV control and prevention.

