## Appendix for "Time-varying transmission dynamics of Novel Coronavirus Pneumonia in China"

### 1.1 Definition and identification of 2019-nCoV case

A suspected case was diagnosed if he/she met one of the following epidemic history and two of the following clinical symptoms. Epidemic history included: (1) a history of travel to or a person who lived in Wuhan or other region where sustained local transmission exists in the 14 days prior to symptom onset; (2) contact with patient with fever/respiratory symptoms from Wuhan or other region where sustained local transmission exists in the 14 days prior to symptom onset; (3) came from a cluster of 2019-nCoV cases or have epidemic relationship with 2019-nCoV cases. Clinical symptoms included: (1) fever; (2) chest radiologic changes (multiple mottling and interstitial change at early phase, bilateral pulmonary multiple ground-glass opacity, infiltrates at advanced phase, lung consolidation at the phase of severe illness); (3) total white blood cell count is normal/low or total lymphocyte count is low. A confirmed case was defined if a suspected case was verified 2019-nCoV by real-time reverse transcriptase polymerase chain reaction assay (RT-PCR) or genetic sequencing.

### 1.2 The association of onset date and reporting date of NCP

A GAM model was employed to estimate the association of onset date and reporting date of NCP cases.

$$E(Y_t) = S(X_{t-i}, k=3) \qquad (1)$$

Where $E(Y_t)$ denotes the expected number of reporting cases on day $t$. $X_{t-i}$ denotes the number of incidences on $i$ days prior to day $t$. For example, the number of reporting cases was 120 on February 1, 2020 in Guangdong Province, in which 10 cases had onset of symptoms on January 31, 20 cases had symptoms on January 30, 35 cases had symptoms on January 29, *etc*. The longest lag was ten days. The association of daily number of reporting cases with daily number of incidences can been seen in Figure S1.

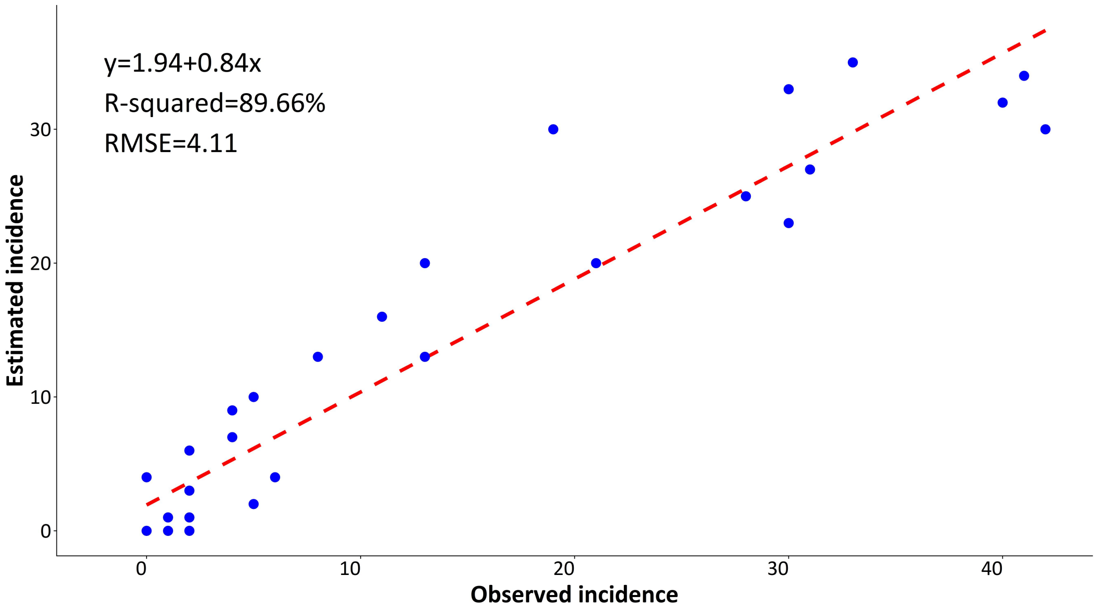


Figure S1: Performance and validation of modeling of predicting the daily incidence in Guangdong Province, China

RMSE: root mean squared prediction error (Cases/day).

Blue dots represent the daily number of NCP cases.

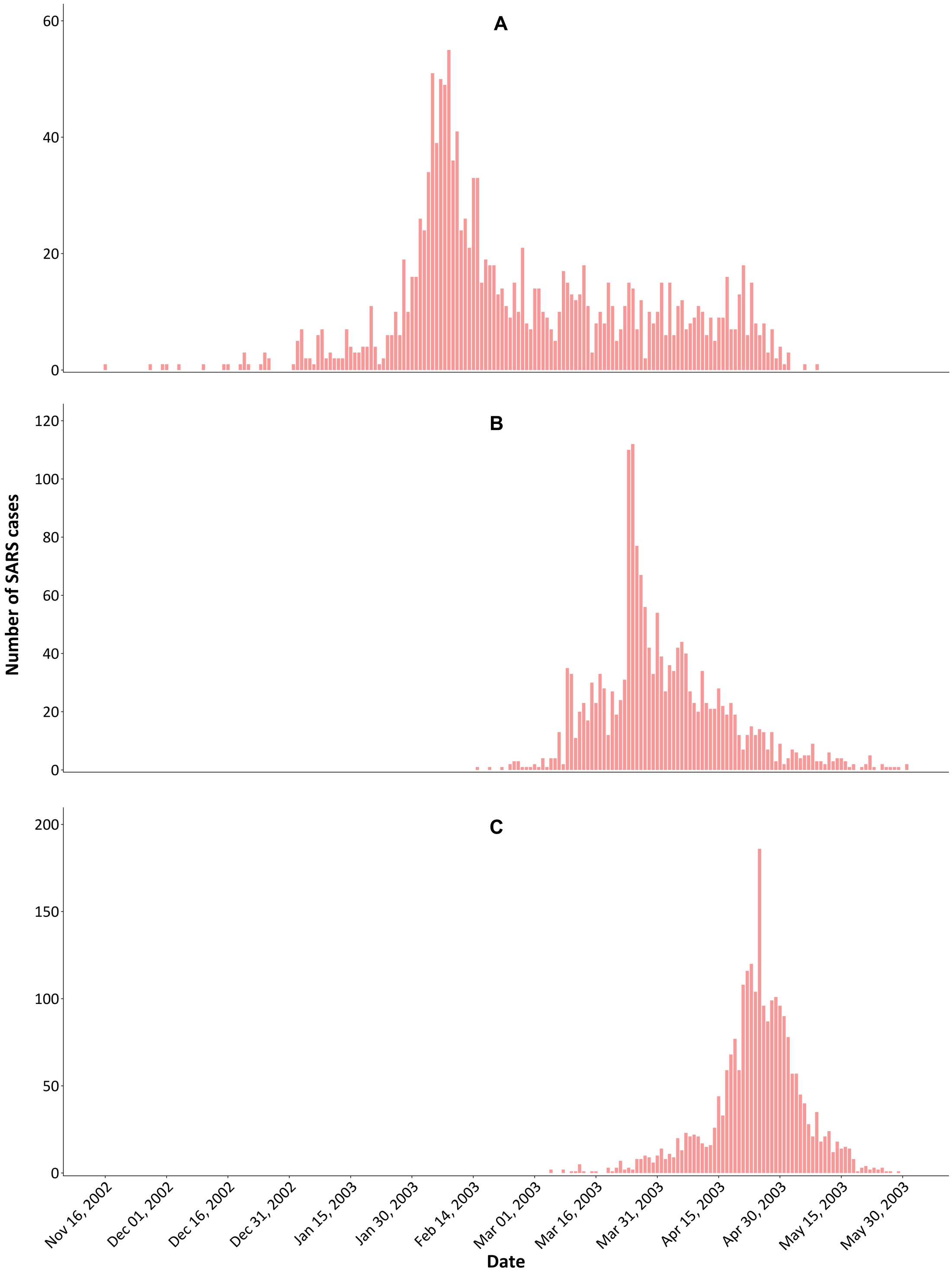


Figure S2. Temporal distribution of confirmed SARS cases in 2002-2003 in China

Panel A: Temporal distribution of SARS cases in Guangdong Province;

Panel B: Temporal distribution of SARS cases in Hong Kong;

Panel C: Temporal distribution of SARS cases in Beijing;

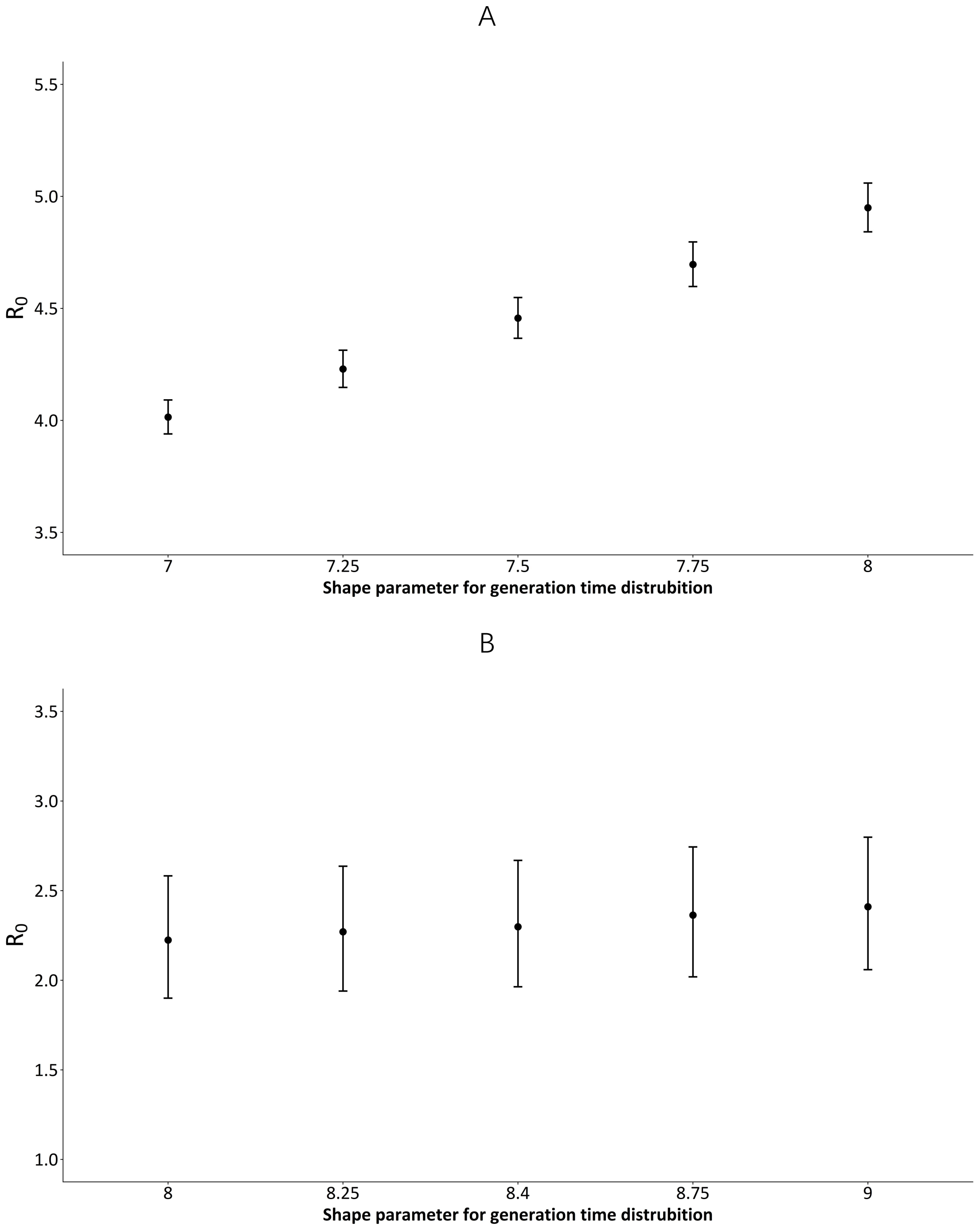


Figure S3. Sensitivity analyses on the impacts of GT on the $R_0$

Panel A: Sensitivity analyses for the NCP cases nationwide;

Panel A: Sensitivity analyses for the SARS cases in Guangdong Province;

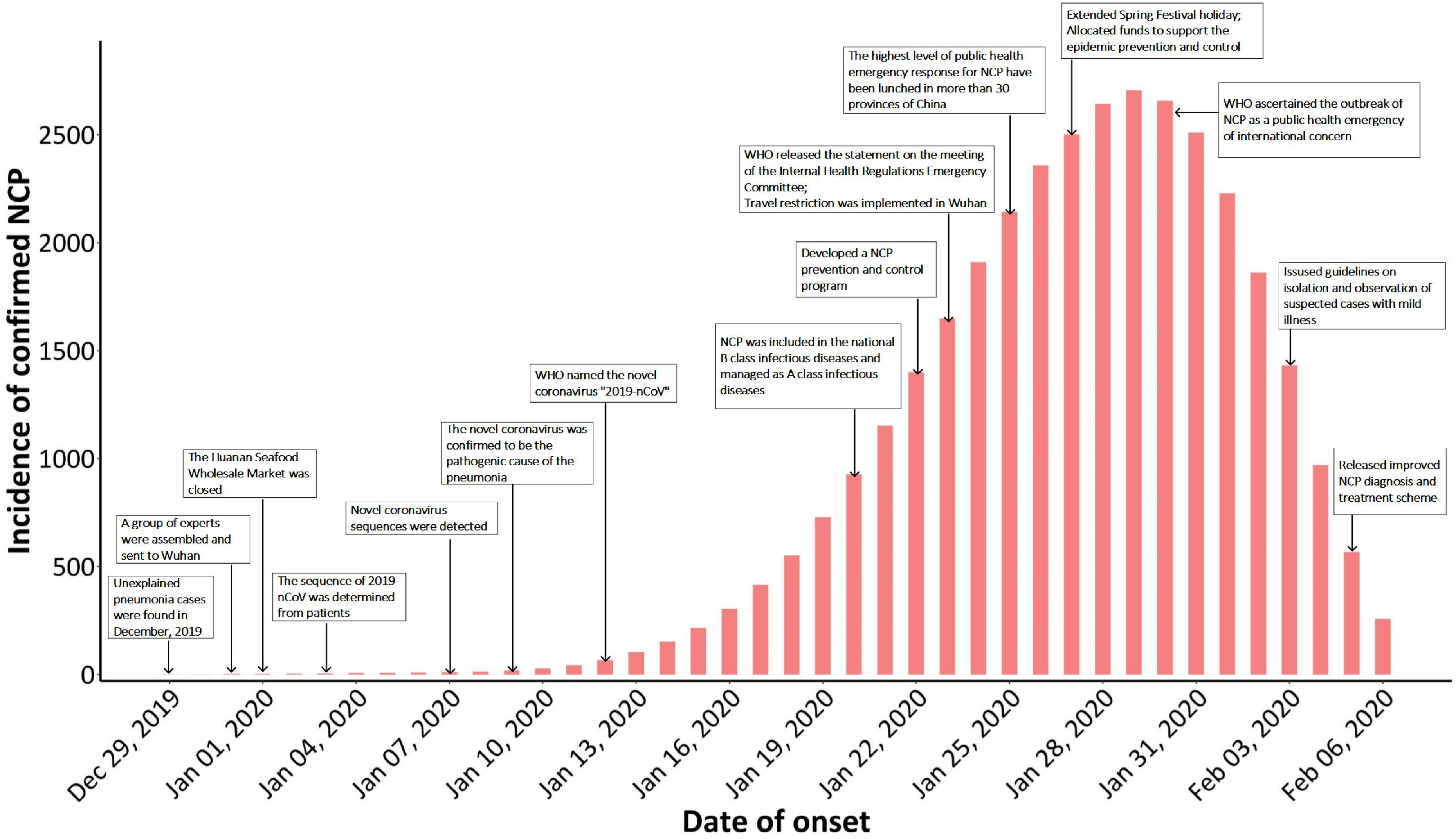


Figure S4. Epidemic curve for 2019-nCoV outbreak and timeline of major control measures up to February 6, 2020 in China

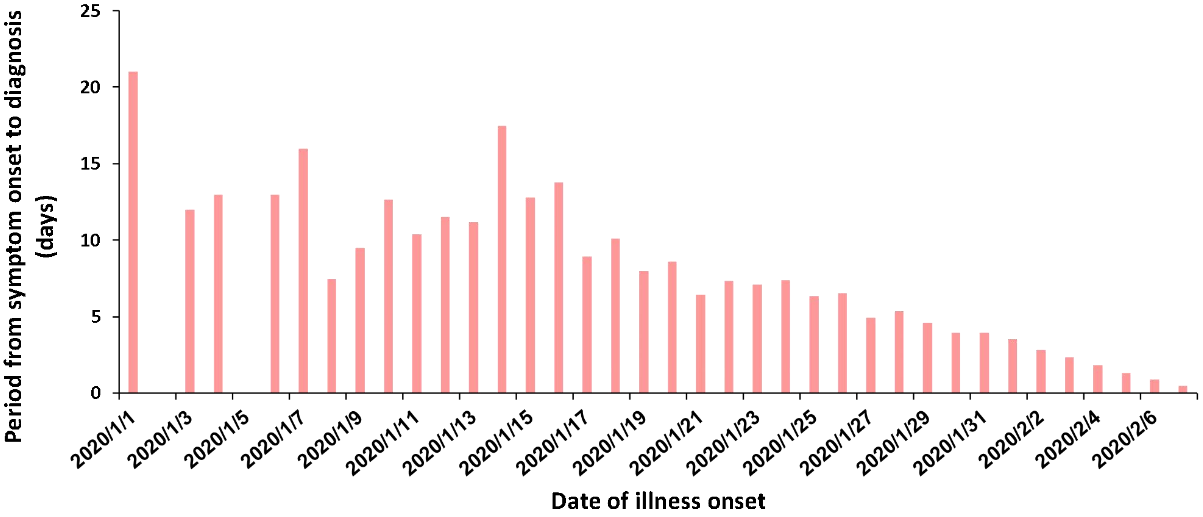


Figure S5. The temporal change of period from symptom onset to diagnosis in NCP cases in Guangdong Province
